# MOD-scTWAS: Leveraging gene co-expression for single-cell transcriptome-wide association studies

**DOI:** 10.64898/2026.09.24.754034

**Authors:** Hanmin Guo, Yao Lu, Lin Hou

**Author notes:** These authors contributed equally to this work (Hanmin Guo, Yao Lu).

## Abstract

Transcriptome-wide association studies (TWAS) provide an effective framework for identifying genes associated with complex traits. Population-scale single-cell transcriptomic data enable genetically regulated expression (GReX) prediction and TWAS analyses at cell-type resolution, but the predictive performance of existing single-cell TWAS methods remains limited. Here, we develop MOD-scTWAS, a module-based method that jointly models GReX for genes within co-expression modules to borrow information across genes. Starting from a generative model for single-cell gene expression, MOD-scTWAS accounts for the heteroscedasticity and cross-gene correlation of individual-level pseudobulk expression in joint GReX prediction. In cross-validation analyses of the OneK1K dataset, MOD-scTWAS achieved higher mean GReX prediction accuracy than scTWAS across all 14 cell types and increased the number of imputable genes. When applied to TWAS analyses of UK Biobank quantitative hematological traits, MOD-scTWAS identified more significant cell type–gene–trait associations than scTWAS. These results demonstrate the potential of leveraging gene co-expression through joint modeling to improve cell-type-specific GReX prediction and TWAS discovery.

## 1 Introduction

Transcriptome-wide association studies (TWAS) integrate eQTL datasets with genome-wide association study (GWAS) data to investigate associations between genetically regulated expression (GReX) and complex traits (Gusev et al., 2016; Gamazon et al., 2015). The predictive performance of GReX models determines the set of genes available for downstream association analysis and influences the statistical power to detect gene–trait associations (Fryett et al., 2020). Improving GReX prediction is therefore an important objective in TWAS methodology. Previous TWAS methods have sought to improve GReX prediction using additional sources of information, including functional annotations (Zhang et al., 2019; Melton et al., 2024) and trans-regulatory information (Luningham et al., 2020; Bhattacharya et al., 2021). Joint modeling across multiple tissues has also been widely used to borrow information for tissue-specific GReX prediction (Hu et al., 2019; Zhou et al., 2020; Song et al., 2024).

Single-cell transcriptomic and eQTL data have enabled TWAS analyses at cell-type resolution (Zeng et al., 2024; Bian et al., 2025; Abe et al., 2025). For example, Zeng et al. (2024) constructed pseudobulk expression profiles from single-cell data and performed TWAS using FUSION (Gusev et al., 2016), identifying gene–trait associations that differed across cell types and subtypes. scTWAS explicitly accounts for the heteroscedasticity of cell-type-specific pseudobulk expression in genetic-effect estimation and improves GReX prediction compared with directly applying conventional TWAS methods to pseudobulk expression (Lin and Su, 2026). In addition, EXPRESSO constructs expression prediction models from cell-type-specific eQTL summary statistics (Wang et al., 2024), and Qin et al. (2026) improve cell-type-specific expression prediction by borrowing information across cell types. scPrediXcan integrates deep learning and single-cell transcriptomic data into a cell-type-specific TWAS framework (Zhou et al., 2025). TWiST and pt-TWAS further extend TWAS to continuous cell states (Qi et al., 2026; Cao et al., 2026).

We propose to leverage gene co-expression to improve GReX prediction in single-cell TWAS. Widespread gene co-expression has been observed across transcriptomic data from different sources and can be used to group genes into distinct co-expression modules, with each module containing genes with similar or related biological functions (Van Dam et al., 2018; Altman et al., 2021). Gene co-expression may arise from multiple mechanisms, including shared upstream regulation, direct or indirect regulatory relationships among genes, and cell states or biological processes that simultaneously affect the expression of multiple genes (Van Dam et al., 2018; Xulvi-Brunet and Li, 2010; Kotliar et al., 2019; Farahbod and Pavlidis, 2020). Existing TWAS methods have leveraged information from co-expressed genes or coexpression modules to capture potential trans-regulatory effects and improve GReX prediction of target genes (Rossi et al., 2026; Brunton et al., 2026). However, joint modeling of multiple co-expressed genes to improve GReX prediction from cis-SNPs has not been explored. We reason that jointly modeling genes within co-expression modules may provide an opportunity to borrow information across genes and improve cell-type-specific GReX prediction.

Motivated by this idea, we develop MOD-scTWAS, a module-based method for singlecell TWAS. Starting from a generative model for single-cell gene expression, MOD-scTWAS derives the heteroscedasticity and cross-gene covariance structure of individual-level pseudobulk expression and uses these results to construct a joint working model for GReX prediction across multiple genes. MOD-scTWAS uses penalized regression to jointly estimate the cis-SNP effects on genes within a module, with the covariance matrix pre-estimated under a working model that ignores genetic regulation. We evaluate the cell-type-specific GReX prediction performance of MOD-scTWAS using the OneK1K single-cell dataset and apply the resulting prediction models to TWAS analyses of UK Biobank quantitative hematological traits. Compared with scTWAS (Lin and Su, 2026), MOD-scTWAS improves mean GReX prediction accuracy, increases the number of imputable genes, and identifies more significant TWAS gene–trait associations. Further analyses suggest that incorporating within-module cross-gene correlation contributes to the improved performance of MOD-scTWAS. We further highlight selected co-expression modules to illustrate potential mechanisms and cell-type contexts underlying gene co-expression, and present additional TWAS discoveries supported by existing functional and genetic evidence.

## 2 Results

### 2.1 MOD-scTWAS Improves Cell-Type-Specific GReX Prediction

We compared the cell-type-specific GReX prediction performance of MOD-scTWAS and scTWAS across 14 peripheral blood cell types in the OneK1K dataset. For each cell type, prediction performance was evaluated using individual-level five-fold cross-validation, with out-of-fold GReX predictions obtained for all analyzed individuals. Weighted predictive *R*^2^ was used as the primary evaluation metric, and genes with weighted predictive *R*^2^ *>* 0.01 were considered imputable. To ensure a direct comparison, the two methods used the same genes, candidate cis-SNPs, individuals, and covariates within each cell type (see Methods for details). Cross-validation results showed that MOD-scTWAS generally achieved higher GReX prediction accuracy across cell types. In all 14 cell types, MOD-scTWAS achieved a higher mean weighted predictive *R*^2^ than scTWAS (Figure 1a); the results were consistent when unweighted predictive *R*^2^ was used for evaluation (Figure S1). At the weighted predictive *R*^2^ *>* 0.01 imputability threshold, MOD-scTWAS yielded more imputable genes in 11 of the 14 cell types, with fewer imputable genes than scTWAS in only CD8_S100B_, DC, and NK_R_ (Figure S2); all three of these cell types had relatively small numbers of cells in the OneK1K dataset. Across the 14 cell types, MOD-scTWAS yielded a total of 8,695 imputable cell type–gene pairs, compared with 8,343 for scTWAS. When different weighted predictive *R*^2^ thresholds were further examined, MOD-scTWAS generally retained an advantage in the number of imputable cell type–gene pairs across the range of thresholds considered (Figure 1b). In addition, GReX prediction accuracy differed substantially across cell types, with cell types containing larger numbers of cells in the OneK1K dataset generally showing higher mean weighted predictive *R*^2^ and more imputable genes.

**Figure 1:**
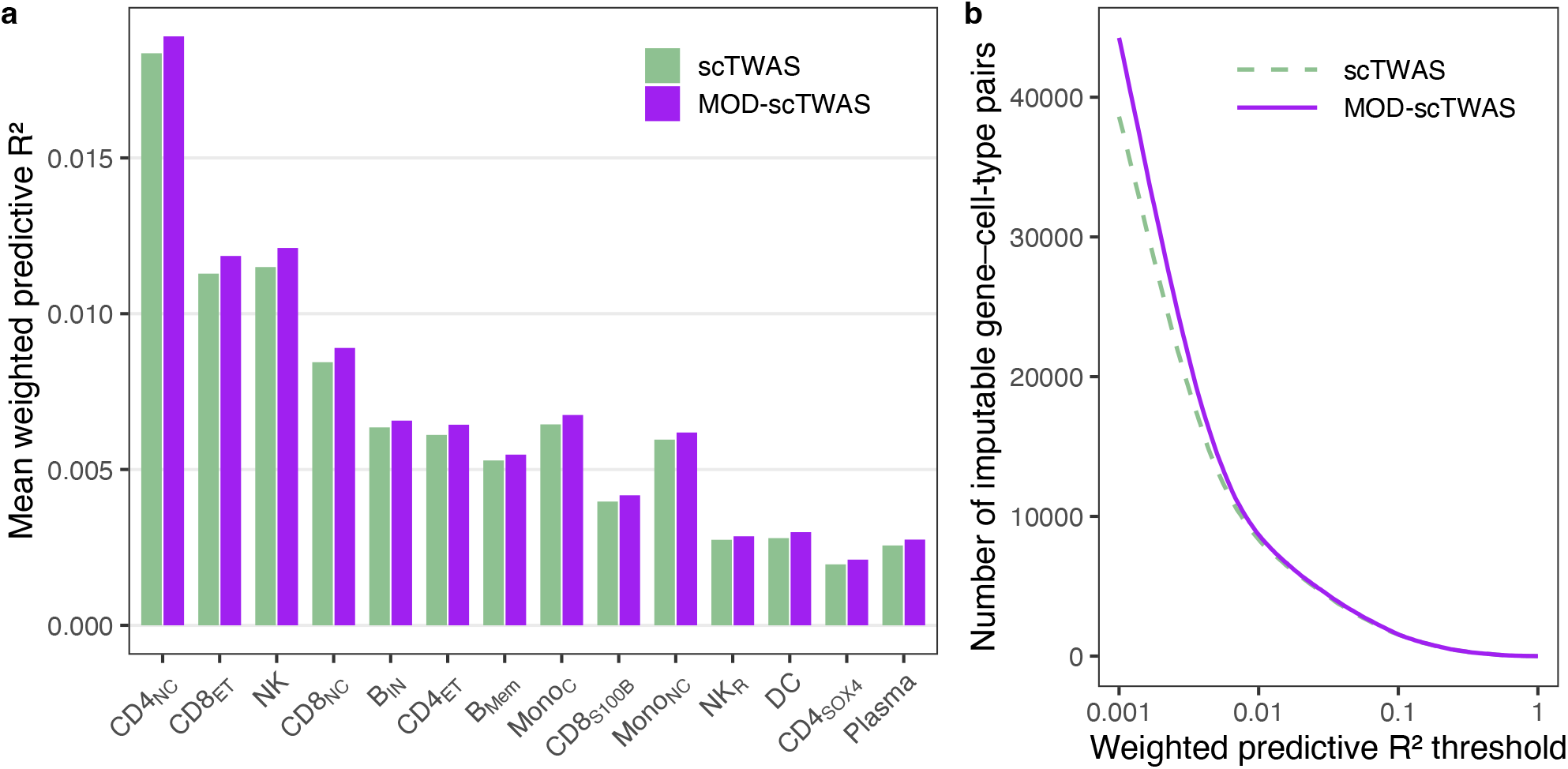
Evaluation of MOD-scTWAS across cell types in the OneK1K dataset. **(a)** Mean weighted predictive *R*^2^ of scTWAS and MOD-scTWAS across 14 peripheral blood cell types. Cell types are ordered by the corresponding number of cells in the OneK1K dataset in descending order. **(b)** Total number of cell type–gene pairs across the 14 cell types reaching each weighted predictive *R*^2^ threshold for the two methods. The x-axis is shown on a logarithmic scale.

### 2.2 Prediction Gains Are Associated with Within-Module Cross-Gene Correlation

The improvement in prediction accuracy achieved by MOD-scTWAS may benefit from its use of gene co-expression information. For target gene *j* in cell type *t*, we defined its within-module cross-gene correlation strength *S*_*jt*_ as the mean of the squared estimated correlations, 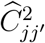, between the target gene and the other genes *j*^*′*^ in the same module; modules containing only one gene after preprocessing were excluded from this analysis. Based on the empirical distribution of *S*_*jt*_ across all cell type–gene pairs included in the analysis, we classified the pairs into low-correlation (0th–50th percentile), moderate-correlation (50th–90th percentile), and high-correlation (90th–100th percentile) groups. The improvement in weighted predictive *R*^2^ of MOD-scTWAS relative to scTWAS generally increased with within-module cross-gene correlation strength, with the largest improvement observed in the high-correlation group (Figure 2a).

**Figure 2:**
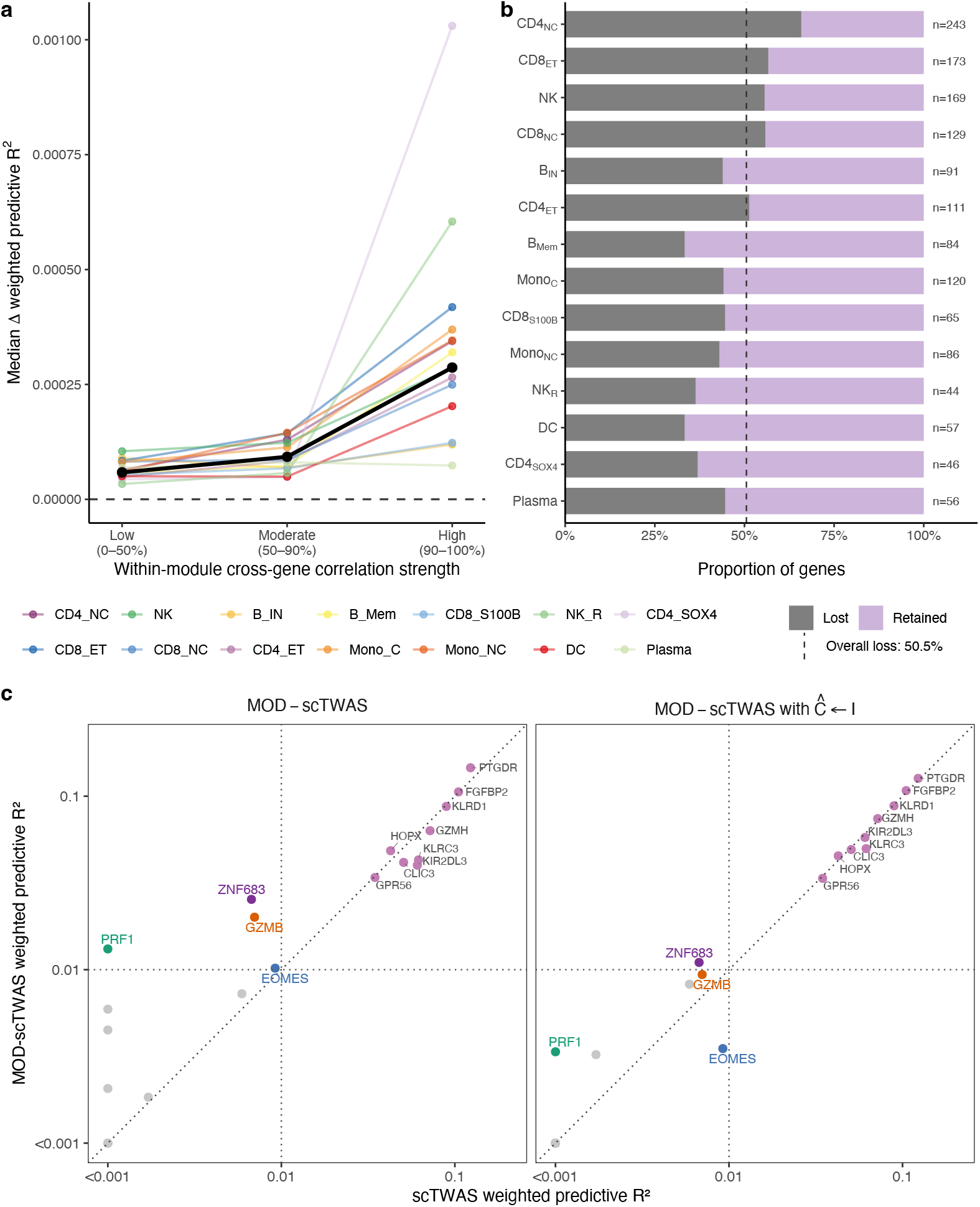
Relationship between cross-gene correlation structure and improvements in GReX prediction performance with MOD-scTWAS. **(a)** Relationship between within-module crossgene correlation strength (*S*_*jt*_) and improvement in GReX prediction performance. Within each of the three correlation-strength groups, the median 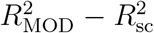 is shown for each cell type, where 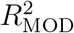 and 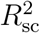 denote the weighted predictive *R*^2^ values from MOD-scTWAS and scTWAS, respectively; different colors represent different cell types, and the black line represents the median across all cell type–gene pairs. **(b)** Ablation analysis of cell type–gene pairs that were imputable only with MOD-scTWAS. Bars show the proportions of pairs that remained imputable or lost imputability after replacing the estimated cross-gene correlation matrix with the identity matrix; the dashed line indicates the overall proportion that lost imputability. **(c)** Example from the BloodGen3 M9.1 module in NK cells. Weighted predictive *R*^2^ values from the full MOD-scTWAS model (left) and the ablation analysis (right) are compared with those from scTWAS. Colored points indicate the four genes discussed in the text; the diagonal dashed line indicates equal predictive *R*^2^ between the two methods, and the horizontal and vertical dashed lines indicate the imputability threshold of *R*^2^ = 0.01.

We further examined the effect of cross-gene correlation structure on GReX prediction performance through ablation analyses in which the cross-gene correlation information was removed from MOD-scTWAS (see Methods for details). We first performed the ablation analysis within modules containing cell type–gene pairs that were imputable only with MOD-scTWAS. Among these MOD-scTWAS-only imputable pairs, 50.5% no longer met the imputability criterion after removing cross-gene correlation information (Figure 2b). We additionally performed the ablation analysis across all analyzed genes in four cell types with different numbers of cells. In each of these cell types, both the mean weighted predictive *R*^2^ and the number of imputable genes were lower than those obtained with the full MOD-scTWAS model (Figure S3). These results support the contribution of cross-gene correlation information to the improved GReX prediction performance of MOD-scTWAS.

Cross-gene correlation structure may arise from different biological mechanisms. For example, BloodGen3 annotates the M10.1 module as an interferon-related module (Altman et al., 2021), and previous studies have shown that type I interferons can induce a shared antiviral transcriptional response across multiple immune cell types (Cui et al., 2024). Imputable genes in this module include the viral RNA-sensing genes *DDX58* and *DHX58*, as well as interferon-response genes such as *IRF7, GBP1, GBP5, PARP14*, and *TRIM22* (Shim et al., 2017; Rehwinkel and Gack, 2020). Within this module, MOD-scTWAS additionally yielded 12 imputable cell type–gene pairs, involving nine genes across six cell types. One example is *SAMD9L*, which was imputable by both methods in NK and CD4_ET_, but only by MOD-scTWAS in CD4_NC_, CD8_ET_, and CD8_NC_; in the ablation analysis, *SAMD9L* no longer met the imputability criterion in any of these three cell types. Previous studies have shown that *SAMD9L* can be induced by interferons across multiple immune cell types (Pappas et al., 2009; Tesi et al., 2017; Legrand et al., 2024). These results suggest that the cross-gene correlation structure leveraged by the joint prediction model in the M10.1 module may arise from an interferon-response transcriptional program broadly present across multiple cell types.

The M13.27 module contains multiple genes involved in T-cell receptor signaling, among which *CD3D, CD247*, and *ZAP70* were classified as imputable only by MOD-scTWAS in some cell types. Both *CD3D* and *CD247* encode components of the T-cell receptor–CD3 complex. Upon receptor activation, phosphorylation of the intracellular immunoreceptor tyrosine-based activation motifs (ITAMs) of CD3/CD247 provides binding sites for ZAP70, which further initiates downstream signaling (Gaud et al., 2018). Previous studies have also observed coexpression of these genes with other genes in the M13.27 module, including *CD6* and *SH2D1A*, in CD8^+^ T cells (Alsulaimany et al., 2022). The coordinated functions of these genes provide a biological basis for their co-expression, and MOD-scTWAS leverages the resulting cross-gene correlation structure to improve GReX prediction.

In NK (natural killer) cells, MOD-scTWAS additionally yielded four imputable genes in the M9.1 module: *EOMES, GZMB, PRF1*, and *ZNF683*. In the ablation analysis, the weighted predictive *R*^2^ decreased for all four genes, three of which no longer met the imputability criterion (Figure 2c). BloodGen3 annotates M9.1 as a cytotoxic lymphocyte-related module, and this module is enriched for natural killer cell-mediated cytotoxicity (Altman et al., 2021). Previous experimental studies further support regulatory relationships among some of the genes in this module. Genetic perturbation and chromatin-binding analyses showed regulatory relationships between EOMES and multiple genes in the M9.1 module, including *PRF1* and *GZMA*, during NK-cell maturation (Zhang et al., 2021). CRISPR perturbation of *EOMES* and *TBX21* in primary human NK cells also altered the expression of multiple genes in the M9.1 module, including *GZMB, NKG7, PRF1*, and *KLRD1* (Wong et al., 2023). In addition, previous studies have shown that *ZNF683* is involved in human NK-cell development and support a regulatory relationship between *ZNF683* and the M9.1 module gene *SH2D1B* (Post et al., 2017; Li et al., 2022). Taken together, these experimental studies provide evidence for regulatory relationships among these genes in NK cells, while MOD-scTWAS leverages the corresponding cross-gene correlation structure in NK cells to improve GReX prediction performance.

Overall, the improvement in GReX prediction with MOD-scTWAS generally increased with the strength of within-module cross-gene correlation and was attenuated in the ablation analysis when cross-gene correlation information was removed. Literature review of representative modules indicated that the within-module gene co-expression leveraged by MOD-scTWAS involves diverse biological mechanisms, and that these mechanisms may either be broadly shared across multiple cell types or exhibit cell-type specificity.

### 2.3 Improved GReX Prediction Alters Downstream TWAS Discoveries

We next examined how differences in GReX prediction between MOD-scTWAS and scTWAS translated into downstream TWAS discoveries. For each method, cell-type-specific GReX prediction models were applied to 29 UK Biobank quantitative hematological traits, with TWAS restricted to genes that met the corresponding cross-validation imputability criterion. TWAS was performed using GWAS summary statistics, with multiple-testing correction applied separately within each trait and cell type (see Methods for details).

Overall, the two methods shared 17,941 significant cell type–gene–trait discoveries, while 4,913 were identified only by MOD-scTWAS and 4,283 only by scTWAS. MOD-scTWAS also identified more significant gene–trait associations and more TWAS-significant genes than scTWAS after merging duplicate discoveries (Table 1).

**Table 1:**
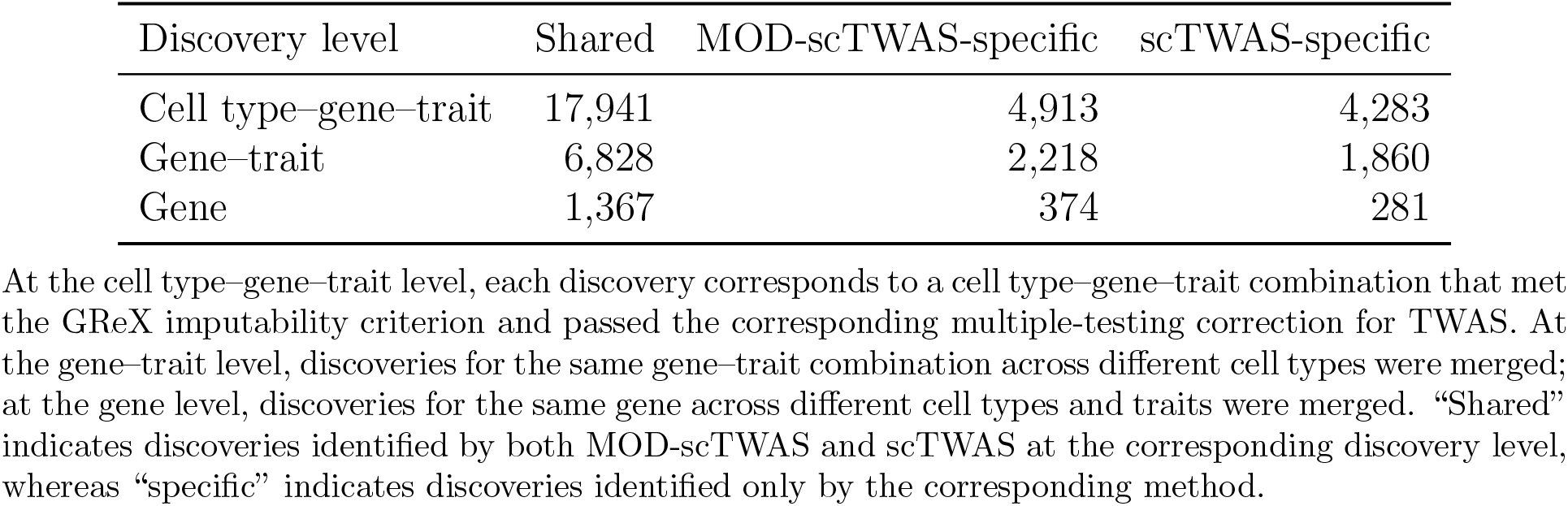
Comparison of TWAS discoveries between MOD-scTWAS and scTWAS.

We grouped TWAS discoveries using the same within-module cross-gene correlation groups defined above and compared the numbers of TWAS discoveries between the two methods within each group. The advantage of MOD-scTWAS in the number of TWAS discoveries was mainly observed in the high-correlation group (Figure 3). Across the 406 trait–cell type combinations, when considering genes in the high-correlation group, MOD-scTWAS yielded more TWAS discoveries in 202 combinations, whereas scTWAS yielded more discoveries in 71 combinations. Aggregated across all 406 combinations, MOD-scTWAS yielded 531 more TWAS discoveries than scTWAS. MOD-scTWAS also showed some advantage in the moderate-correlation group, whereas the numbers of TWAS discoveries were generally similar between the two methods in the low-correlation group. This pattern was consistent with the results from the GReX prediction stage, in which the advantage of MOD-scTWAS was more pronounced among genes with stronger within-module cross-gene correlation.

**Figure 3:**
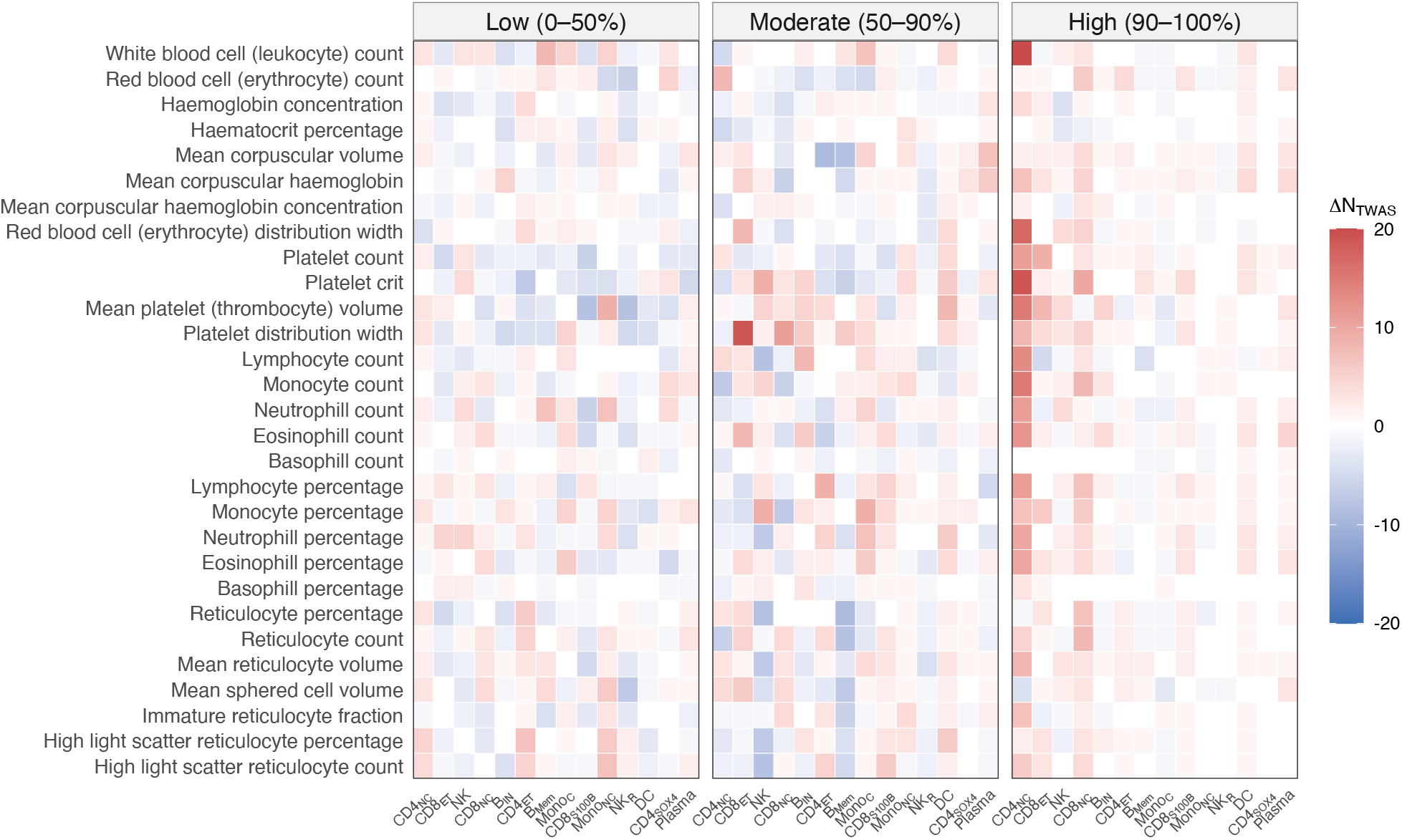
Comparison of the numbers of TWAS discoveries between MOD-scTWAS and scT-WAS across different levels of within-module cross-gene correlation strength. The grouping of cell type–gene pairs by within-module cross-gene correlation strength is the same as in Figure 2a. Heatmap colors represent the number of TWAS discoveries from MOD-scTWAS minus that from scTWAS (Δ*N*_TWAS_) for each hematological quantitative trait and cell type within the corresponding correlation-strength group.

MOD-scTWAS identified several genes that may be relevant to the genetic basis of blood cell traits. We first considered examples from two monocyte subtypes. In Mono_C_ (classical monocytes), the weighted predictive *R*^2^ of *CCR2* increased from 0.0029 with scTWAS to 0.0103 with MOD-scTWAS, thereby reaching the imputability criterion, and *CCR2* showed significant TWAS associations with both monocyte count and monocyte percentage. CCR2 has a well-established functional connection to classical monocytes: previous studies have shown that CCR2 is involved in the egress of classical monocytes from the bone marrow into the peripheral circulation; in human peripheral blood, classical CD14^++^ monocytes express CCR2, whereas CD14^+^CD16^+^ monocytes show substantially lower CCR2 expression (Serbina and Pamer, 2006; Weber et al., 2000). Similarly, in Mono_NC_ (nonclassical monocytes), the weighted predictive *R*^2^ of *CSF1R* increased from 0.0098 to 0.0138, and *CSF1R* showed significant TWAS associations with both monocyte traits described above. Previous studies have shown that CSF1R signaling is involved in the differentiation and homeostasis of nonclassical monocytes; in mice, blockade of CSF1R signaling or knockout of *CSF1R* leads to a selective reduction in Ly6C^low^ nonclassical monocytes (Gallerand et al., 2026). The significant TWAS associations of *CCR2* and *CSF1R* in classical and nonclassical monocytes, respectively, together with existing functional evidence in the corresponding monocyte subtypes, illustrate the potential value of cell-type-specific TWAS analyses for revealing the cell-type-specific genetic basis of complex traits.

In the earlier analysis of GReX prediction performance, we discussed *EOMES* in the M9.1 module in NK cells. Its weighted predictive *R*^2^ increased from 0.0092 with scTWAS to 0.0102 with MOD-scTWAS, just reaching the imputability criterion of *R*^2^ *>* 0.01. Because *EOMES* did not satisfy the scTWAS imputability criterion, it was not included in the formal scTWAS discovery analysis. For comparison, we additionally performed TWAS using the scTWAS prediction weights for *EOMES*. MOD-scTWAS yielded substantially smaller TWAS *p* values for both lymphocyte count and lymphocyte percentage, at 1.20 × 10^−17^ and 1.01 × 10^−11^, respectively, compared with 8.03 × 10^−9^ and 1.42 × 10^−6^ using the scTWAS prediction weights. In contrast, some TWAS discoveries specific to MOD-scTWAS corresponded to larger improvements in GReX prediction accuracy. For example, in B_Mem_, the weighted predictive *R*^2^ of *DTX1* increased from 9.91 × 10^−5^ with scTWAS to 0.0109 with MOD-scTWAS, and *DTX1* was significantly associated with multiple hematological traits, including lymphocyte count with a TWAS *p* value of 1.62 × 10^−8^. Previous experimental studies also support the roles of these two genes in the corresponding cell lineages: EOMES promotes the differentiation and early maturation of human NK cells and contributes to maintaining the transcriptional program and cellular identity of mature NK cells (Kiekens et al., 2021; Wong et al., 2023), whereas DTX1 can antagonize Notch1 signaling and influence the differentiation of lymphoid progenitors toward the B-cell lineage (Izon et al., 2002).

More broadly, the new discoveries from MOD-scTWAS spanned multiple cell types and blood cell traits. In CD4_NC_, *PACSIN2* was significantly associated with multiple platelet-related traits. Previous studies have shown that *PACSIN2* is highly expressed in platelets and is involved in platelet formation; population genetic studies have linked variants in *PACSIN2* to multiple platelet-related traits, and animal experiments have further supported the effects of PACSIN2 on platelet number and function (Begonja et al., 2015; Biswas et al., 2023). Also in CD4_NC_, *ICOSLG* was significantly associated with eosinophil count and eosinophil percentage, and previous large-scale GWAS studies have linked genetic variants near *ICOSLG* to these two eosinophil-related traits (Astle et al., 2016; Vuckovic et al., 2020). *KLRC1* and *HLA-E* were significantly associated with multiple traits, including lymphocyte count, in CD8_ET_ and CD8_NC_, respectively. In CD8^+^ T cells, these two genes participate in the same inhibitory signaling axis: *KLRC1* encodes a component of the inhibitory receptor NKG2A, whereas HLA-E is a ligand for NKG2A; HLA-E–NKG2A signaling can regulate the effector functions of NK cells and CD8^+^ T cells (Pereira et al., 2019).

## 3 Discussion

We developed MOD-scTWAS, which jointly models GReX for multiple genes within coexpression modules to borrow information across genes and improve cell-type-specific GReX prediction and downstream TWAS analyses. Across 14 cell types in the OneK1K dataset, MOD-scTWAS achieved higher mean weighted predictive *R*^2^ than scTWAS in all cell types, increased the total number of imputable cell type–gene pairs from 8,343 to 8,695, and increased the number of imputable genes in 11 cell types. Further analyses suggested that the improvement in GReX prediction performance was closely related to the use of gene co-expression information by MOD-scTWAS: genes with stronger co-expression with other genes within the same module generally showed larger improvements in prediction performance, and when cross-gene correlation was ignored in the joint prediction model by replacing the cross-gene correlation matrix **C** with the identity matrix, approximately 50.5% of the MOD-scTWAS-specific imputable genes no longer met the imputability criterion. In TWAS analyses of UK Biobank quantitative hematological traits, MOD-scTWAS specifically identified 4,913 significant cell type–gene–trait combinations, 630 more than the number of discoveries specific to scTWAS. Further investigation of selected modules provided examples of potential biological sources of the cross-gene correlation structure used by MOD-scTWAS, including transcriptional responses shared across cell types, coordinated expression among genes involved in common biological processes, and gene regulatory relationships within specific cell types. Among the additional TWAS discoveries, *CCR2* and *CSF1R* were significantly associated with monocyte-related traits in classical and nonclassical monocytes, respectively, consistent with previous functional studies in the corresponding monocyte subtypes. Additional associations supported by existing functional or genetic evidence were also observed across multiple immune cell types and hematological traits.

MOD-scTWAS has several limitations. First, MOD-scTWAS requires estimation of multiple types of model parameters, including gene-specific variance parameters and within-module cross-gene correlation matrices. We currently use a stepwise estimation procedure to reduce the complexity of parameter estimation, but have not systematically compared the performance of stepwise and joint estimation. The effects of different estimation strategies on model performance therefore warrant further investigation. Second, the performance of MOD-scTWAS may depend on the total number of cells available for each cell type, particularly because the method requires estimation of within-module cross-gene correlation matrices in addition to the parameters required by single-gene models. Its applicability to rare cell types may therefore be limited. Finally, the BloodGen3 modules used in the current analysis were derived from bulk transcriptomic data and are not cell-type-specific (Altman et al., 2021). Future work could consider co-expression modules constructed from single-cell data or incorporate external information such as sets of downstream response genes identified through CRISPR perturbation experiments to further optimize the gene sets used for joint modeling.

## 4 Methods

### 4.1 Generative Model for Single-Cell and Pseudobulk Gene Expression

Consider single-cell transcriptomic data from a given cell type. For the *c*th cell from individual *i*, we jointly consider *G* genes across the genome, where *i* = 1, …, *n* and *c* = 1, …, *n*_*i*_. We model the latent gene expression levels, the single-cell transcription process, and the RNA capture and sequencing process in sequence, and derive the resulting variance and cross-gene covariance structure of individual-level pseudobulk expression.

First, let the latent expression levels of the *G* genes in the *c*th cell from individual *i* be

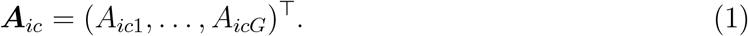

For the *j*th gene, we consider the following additive model:

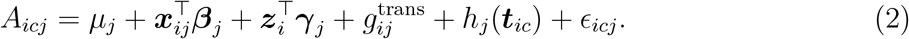

We assume that the probability that the additive model above takes negative values is negligible. Here, *µ*_*j*_ denotes the baseline expression level of gene *j*; ***x***_*ij*_ is the column vector of cis-SNP genotypes near gene *j* for individual *i*, and ***β***_*j*_ denotes the corresponding cis-SNP genetic effects; ***z***_*i*_ is the column vector of individual-level covariates, and ***γ***_*j*_ denotes their corresponding effects; 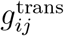 represents the effects of other individual-level genetic regulatory factors, such as trans-SNPs, on the expression of gene *j*; ***t***_*ic*_ is a random vector of cell states, and *h*_*j*_(·) is a fixed multivariate function such that *h*_*j*_(***t***_*ic*_) represents variation in gene expression attributable to changes in cell state; and *ϵ*_*icj*_ represents other unexplained cell-level random effects.

In this model, *µ*_*j*_, ***β***_*j*_, and ***γ***_*j*_ are treated as fixed effects, whereas 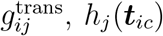, *h*_*j*_(***t***_*ic*_), and *ϵ*_*icj*_ are treated as zero-mean random effects. We assume independence across individuals while allowing these random effects to be correlated across genes. For notational convenience, define the random component of the latent expression level as

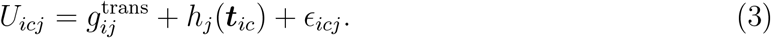

We further assume that *U*_*icj*_ has the same marginal distribution across cells from the same individual, with an exchangeable covariance structure. To fix the scale of the latent expression levels, we impose the constraint

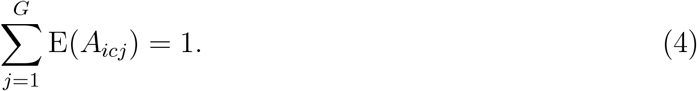

The model above allows the latent expression levels of different genes to be correlated before transcriptional stochasticity and sequencing are introduced. For example, different genes may be influenced by correlated individual-level genetic regulatory factors, while shared cell states or biological processes may induce coordinated changes in the expression of multiple genes. These shared or correlated random factors can therefore induce cross-gene correlation at the level of latent expression.

Conditional on the latent expression levels, we next account for stochasticity in the singlecell transcription process. Motivated by the telegraph model of gene transcription and its steady-state distribution under transcriptional bursting, we use a negative binomial distribution to model the number of RNA molecules within a cell (Shahrezaei and Swain, 2008). Specifically, conditional on the latent expression level of gene *j* in the *c*th cell from individual *i*, we assume

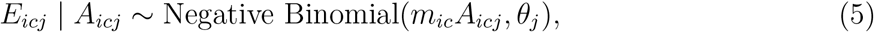

where *m*_*ic*_ denotes the expected total number of RNA molecules in the cell, and *θ*_*j*_ is the dispersion parameter for gene *j*. It follows that

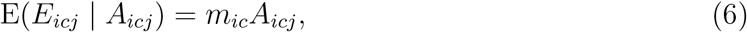

and

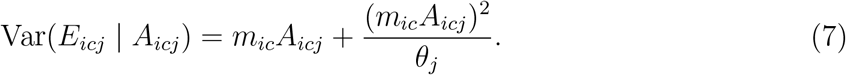

The second term in the variance captures overdispersion relative to a Poisson distribution.

Previous studies have shown that transcriptional bursting across different genes or regulatory elements can exhibit coordinated behavior under specific regulatory mechanisms or biological conditions (Stavreva et al., 2019; Robles-Rebollo et al., 2022). We therefore allow the transcriptional processes of different genes to be correlated. For two distinct genes *j* and *j*^*′*^, we introduce a parameter *r*_*jj′*_ to characterize the degree of correlation, with |*r*_*jj′*_| ≤ 1, and define

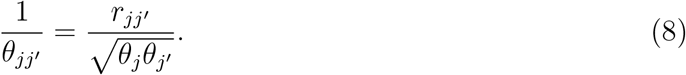

Conditional on the latent expression levels of the two genes, we assume that their RNA molecule counts satisfy

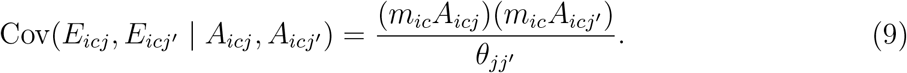

This parameterization provides a second-moment representation of cross-gene dependence arising during the transcription process, motivated by multivariate Gamma–Poisson mixture models (Mosimann, 1963).

Finally, we consider RNA capture and sequencing, followed by the construction of individuallevel pseudobulk expression. Let 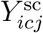 denote the observed single-cell UMI count. Conditional on the actual number of RNA molecules, we assume that the capture and sequencing of different RNA molecules are conditionally independent and that an RNA molecule from individual *i* is captured and sequenced with probability *p*_*i*_. Then,

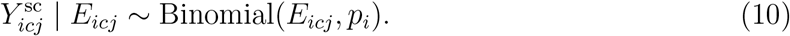

For individual *i* with *n*_*i*_ cells, we define the raw pseudobulk expression of gene *j* as the sum of the corresponding single-cell UMI counts:

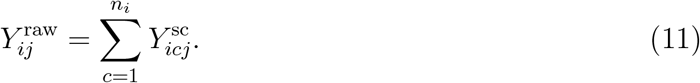

Let *s*_*i*_ denote the total UMI count across all cells and genes for individual *i*. The normalized pseudobulk expression is defined as

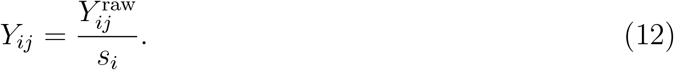

Treating the total UMI count *s*_*i*_ as fixed, the law of total expectation under the hierarchical generative model above gives the expected normalized pseudobulk expression as

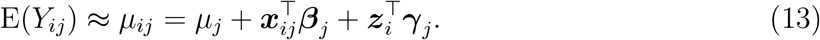

We further assume that *m*_*ic*_ ≈ *m*_*i*_, such that the expected total number of RNA molecules is approximately the same across cells from the same individual. By the law of total variance, the variance of the normalized pseudobulk expression is

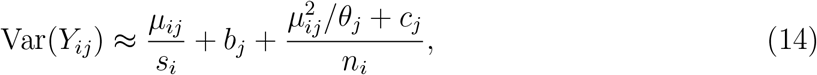

where

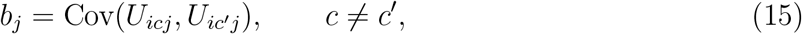

and

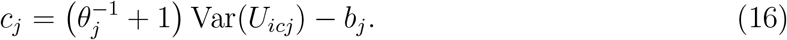

The variance consists of three components. The first depends on the total UMI count *s*_*i*_ and reflects the effect of sequencing depth. The second, *b*_*j*_, reflects the contribution of individual-level random effects. The third captures the combined contribution of cell-level random effects and transcriptional overdispersion, and decreases as the number of cells *n*_*i*_ increases. Consequently, differences in total UMI counts and cell numbers across individuals induce heteroscedasticity in pseudobulk expression.

Similarly, for two distinct genes *j* and *j*^*′*^, the covariance between their normalized pseudobulk expression levels is

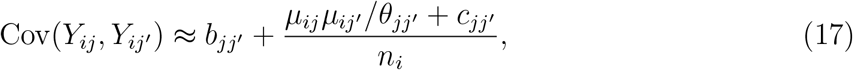

where

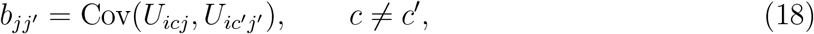

and

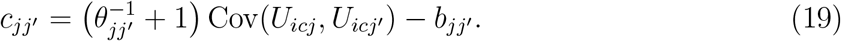

The covariance has a form analogous to that of the variance and arises from cross-gene correlations in individual-level and cell-level random effects, as well as correlations between the transcriptional processes of different genes.

Taken together, the hierarchical generative model shows that aggregating single-cell expression data to the individual level gives rise to both heteroscedasticity and cross-gene covariance in pseudobulk expression. When leveraging cross-gene correlation to improve GReX prediction in the subsequent working model, we do not further distinguish among the specific biological sources underlying these correlations.

### 4.2 Multivariate Working Model for Joint GReX Prediction

To jointly predict GReX for multiple genes within a gene module, we construct a working model based on the generative model for single-cell gene expression described above. For a gene module containing *p* genes, let

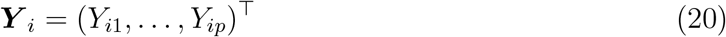

denote the vector of normalized pseudobulk expression levels for individual *i*. The variance and covariance structures derived in the previous subsection (Equations (14) and (17)) involve multiple gene-specific and gene-pair-specific parameters. To reduce model complexity while retaining gene-specific heteroscedasticity and cross-gene correlation, we adopt the following simplified working covariance structure:

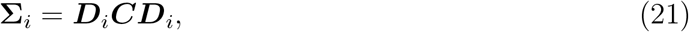

where

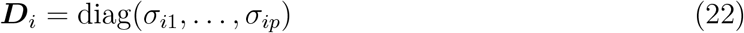

is a diagonal matrix containing the gene-specific expression standard deviations for individual *i*, and ***C*** is the within-module cross-gene correlation matrix shared across individuals. Further, we simplify the term 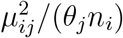 in Equation (14) by ignoring its between-individual variation arising from *µ*_*ij*_, and reparameterize the corresponding gene-specific component using *a*_*j*_ as

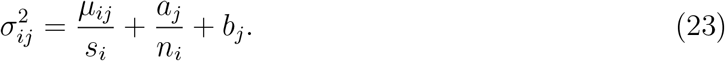

Based on the covariance structure above, we specify the following working model:

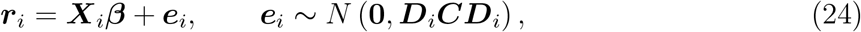

where ***r***_*i*_, ***X***_*i*_, and ***β*** denote the vector of gene expression residuals, the cis-SNP design matrix, and the vector of cis-SNP genetic effects within the module, respectively. Specifically,

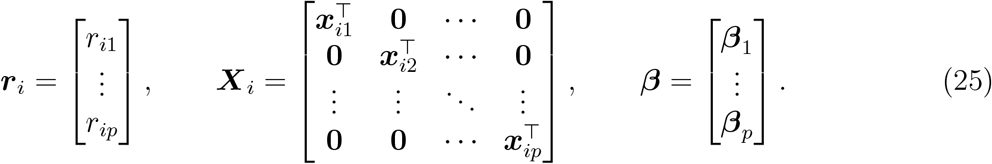

Here, *r*_*ij*_ denotes the expression residual for gene *j* after removing the mean and the effects of individual-level covariates; its calculation is described in Section 4.3. The vector ***x***_*ij*_ contains the standardized cis-SNP genotypes for gene *j*, and ***β***_*j*_ denotes the corresponding cis-SNP genetic effects.

### 4.3 Estimation of Marginal Variance and Cross-Gene Correlation

To calculate the expression residuals and estimate the covariance matrices required for the joint working model, we first consider the null model with no cis-genetic effects, corresponding to ***β*** = **0** in Equation (24). For gene *j*, the corresponding marginal working model is then

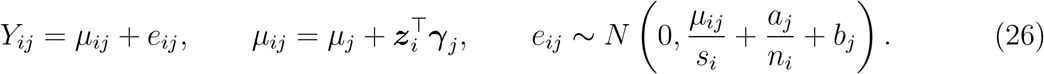

To estimate the mean parameters, *µ*_*j*_ and ***γ***_*j*_, together with the variance parameters, *a*_*j*_ and *b*_*j*_, we use a one-step updating procedure similar to iteratively reweighted least squares (IRLS). We first set *a*_*j*_ and ***γ***_*j*_ to zero and initialize the mean parameter *µ*_*j*_ using the sample mean. Consequently, the initial fitted mean for each individual is

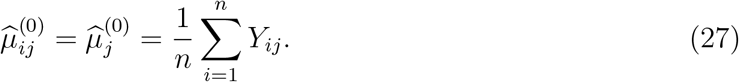

Holding 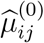 fixed, we estimate the variance parameters using the method-of-moments procedure described below, yielding 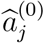 and 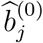. The initial variance estimate is then

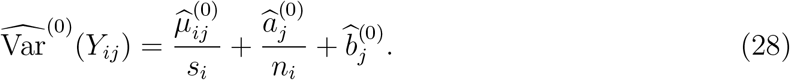

For the one-step update, we perform weighted linear regression using the inverse of 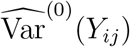 as weights to obtain updated estimates of the mean parameters, 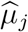 and 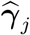. Let

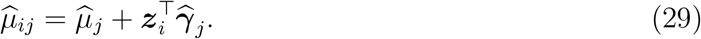

Holding the updated 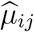 fixed, we re-estimate *a*_*j*_ and *b*_*j*_ using the same method-of-moments procedure, yielding the final marginal variance estimate

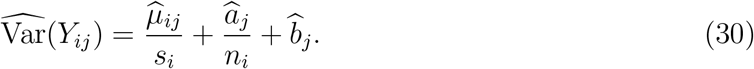

The variance parameters *a*_*j*_ and *b*_*j*_ are estimated by the following method-of-moments procedure. Let 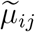 denote the current estimate of the mean, corresponding to either 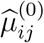 or 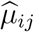. From Equation (26),

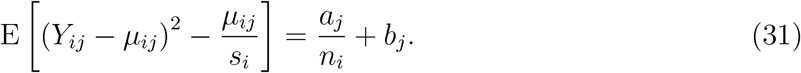

Plugging 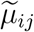 in for *µ*_*ij*_, we construct the following two moment equations:

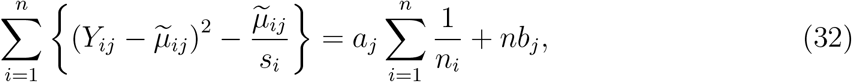

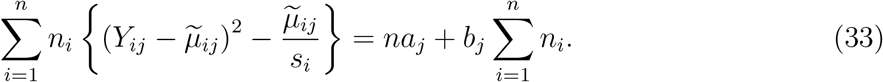

Solving these two equations simultaneously yields the method-of-moments estimates of *a*_*j*_ and *b*_*j*_, with negative estimates truncated at zero.

After fitting the marginal model separately for each gene within a module, we define the expression residuals as

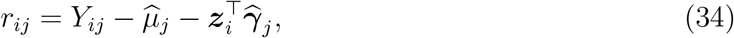

and combine them across genes within the module to form the residual vector in the joint working model,

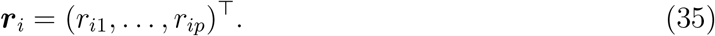

Using the final marginal variance estimates for each gene, we construct

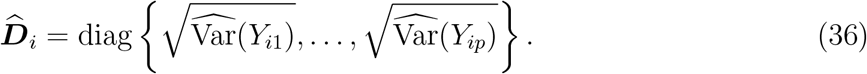

Under the joint null model described above, we estimate the cross-gene correlation matrix ***C*** by standardizing the expression residuals using 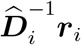 and calculating the sample correlation matrix of the standardized residuals across individuals within the module. We denote the resulting estimate by 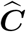. The estimated covariance matrix for individual *i* is then

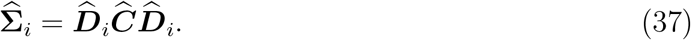

### 4.4 Joint Penalized Estimation of GReX Prediction Models

We estimate the model (24) using penalized generalized least squares with an elastic net penalty:

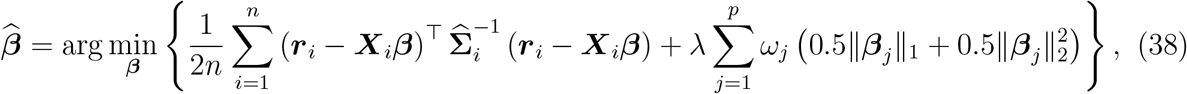

where *λ* is a tuning parameter controlling the overall penalty strength, and *ω*_*j*_ is a gene-specific penalty factor that adjusts the relative penalty strength across genes (see Section 4.5).

Given 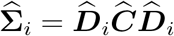, let 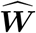 be any matrix satisfying

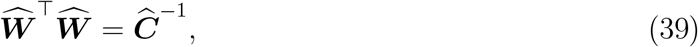

and define

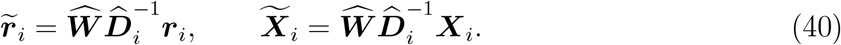

The optimization problem in Equation (38) can then be equivalently written as

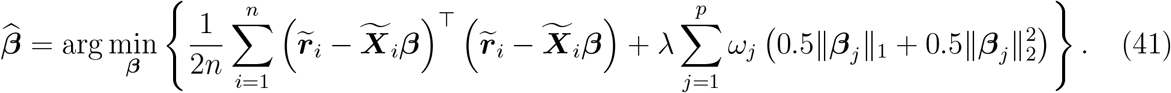

This optimization problem can be solved using standard elastic net algorithms. We fit the model using the R package `glmnet`, with *λ* selected by 10-fold cross-validation within the training data based on mean squared prediction error.

This procedure yields the joint estimates of the cis-SNP genetic effect vectors for genes within the module,

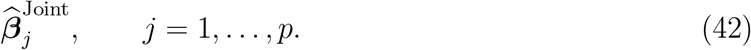

For a new individual *i*^*\**^, let 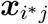 denote the genotype vector of the corresponding cis-SNPs for gene *j*. The predicted GReX is then

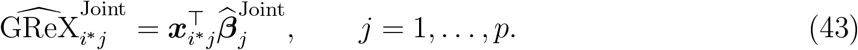

Because the tuning parameter *λ* is selected based on cross-validated prediction performance across multiple genes within a module, the resulting penalty strength may be suboptimal for individual genes. For any gene with 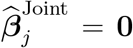, we therefore refit the prediction model for gene *j* alone, with *λ* reselected using the same 10-fold cross-validation procedure. We denote the resulting single-gene GReX prediction for the new individual *i*^*\**^ by 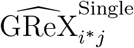 and define the final MOD-scTWAS GReX prediction as

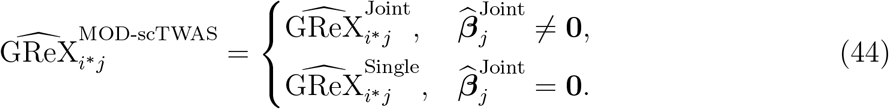

### 4.5 Practical Strategies for Joint Model Fitting

#### Gene module refinement

The definition of gene modules determines the set of genes among which information is borrowed in the joint prediction model. Information borrowing arises from the equivalent transformation in Equation (41), through which the cross-gene correlation structure induces mixing of expression residuals and predictors across genes. We therefore use previously defined gene co-expression modules as the initial modules and adaptively refine them based on the gene correlation matrix 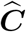 estimated from the training data. The refinement is designed to limit the extent of information borrowing across genes, ensure numerical stability when inverting the correlation matrix, and control the computational scale of joint model fitting.

First, let

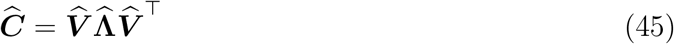

denote the eigendecomposition of 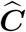, and define the symmetric whitening transformation

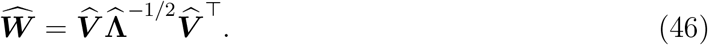

To separately control the information borrowed by gene *j* from all other genes and from any single other gene, we define

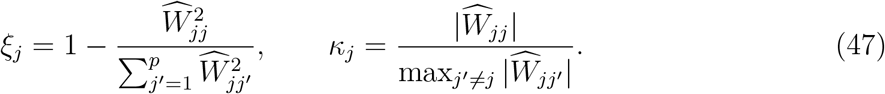

To assess numerical stability of matrix inversion, we use the minimum eigenvalue of 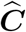 as a diagnostic criterion. In addition, to control the computational scale of multigene joint fitting, we restrict the number of genes within each module. Based on these considerations, a module is no longer refined if it simultaneously satisfies

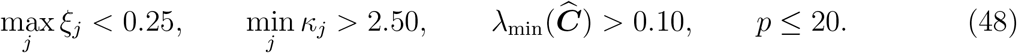

Otherwise, the module is further subdivided.

To preserve the within-module gene correlation structure during refinement, we partition modules using ideas from Fiedler graph partitioning and normalized cut (Ncut) (Fiedler, 1973; Shi and Malik, 2000). For a module requiring further subdivision, we construct a nonnegative symmetric adjacency matrix ***A*** from the positive correlations in its estimated correlation matrix 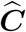:

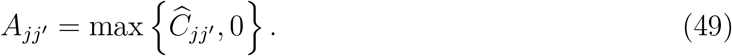

We then construct the graph Laplacian

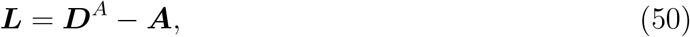

where ***D***^*A*^ is the degree matrix whose diagonal elements are the row sums of ***A***. We perform an eigendecomposition of ***L***, obtain the Fiedler vector ***v***_2_ corresponding to its second-smallest eigenvalue, and order the genes within the module according to the elements of ***v***_2_. We evaluate all possible split points along this ordering to partition the genes into two subsets and select the bipartition with the minimum Ncut value. The assessment and refinement procedure is recursively applied to each resulting submodule until the stopping criteria are satisfied.

#### Gene-specific penalty factors

Because expression variances differ across genes, the loss function in Equation (41) assigns different weights to different genes. Consequently, when the overall penalty parameter *λ* is shared across genes within a module, the relative penalty strength may differ across genes. We therefore adjust the gene-specific penalty strength by considering the Karush–Kuhn–Tucker (KKT) conditions of the joint prediction model under ***C*** = **I**. An approximate analysis of the KKT conditions shows that whether all coefficients for gene *j* are shrunk to zero depends on the quantity

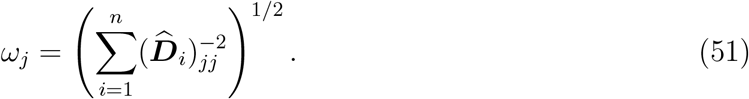

Therefore, to reduce the influence of gene-specific expression variances on the relative penalty strength, we use *ω*_*j*_ as the gene-specific penalty factor in Equation (38).

#### SNP prescreening

To reduce the computational burden arising from the large number of cis-SNPs in multigene joint models, we prescreen candidate cis-SNPs for each gene before model fitting. This strategy follows the idea of sure independence screening (SIS) (Fan and Lv, 2008), in which candidate predictors are screened according to their marginal correlations with the response.

To account for the heteroscedasticity of pseudobulk expression, we calculate both the sample correlation between SNP genotypes and the gene expression residuals *r*_*ij*_ defined in Equation (34), and a weighted sample correlation using the inverse of the variance estimate in Equation (30) as weights. For gene *j* with *m*_*j*_ candidate cis-SNPs, we define

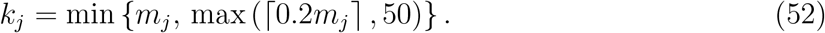

We rank the cis-SNPs separately by the absolute values of the two correlation measures and retain the top *k*_*j*_ cis-SNPs under each ranking. The union of the two sets of retained SNPs is then used as the predictor set for gene *j* in the joint prediction model.

### 4.6 OneK1K Data and Analysis Procedures

We applied MOD-scTWAS and scTWAS (Lin and Su, 2026) to the population-scale single-cell eQTL dataset OneK1K (Yazar et al., 2022). Following the recommendations of the original study authors, a total of 980 individuals were included in the analysis. Analyses were conducted separately for 14 peripheral blood cell types. To ensure a fair comparison between the two methods, MOD-scTWAS and scTWAS used the same genes, initial candidate cis-SNP pools, individuals, and covariates within each cell type. Following the pseudobulk definition above, we summed the raw UMI counts across all cells of the same cell type from each individual on a gene-by-gene basis to obtain *Y* ^raw^, and normalized these counts to obtain 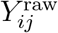. Individual-level covariates included sex, age, and the first six genotype principal components used in the OneK1K study. For each cell type, we analyzed genes that were included in the original OneK1K eQTL analysis and were also covered by the BloodGen3 transcriptional modules (Altman et al., 2021), resulting in 4,790–7,848 genes across cell types. For each retained gene, both methods used the corresponding cis-SNPs from the original OneK1K eQTL analysis as candidate genetic predictors. For MOD-scTWAS, genes appearing in multiple BloodGen3 modules were assigned to the module with the smallest size.

### 4.7 Evaluation of GReX Prediction

We evaluated GReX prediction accuracy using five-fold cross-validation at the individual level. For each cell type, individuals were randomly divided into five folds. For each fold, both methods estimated GReX prediction coefficients using four-fifths of the individuals as the training set and predicted GReX for individuals in the held-out fold. This procedure was repeated across the five folds to obtain out-of-fold GReX predictions for all individuals. All model fitting and computational procedures described above were performed using only the corresponding training set.

In TWAS studies, GReX prediction accuracy is commonly evaluated using predictive *R*^2^, defined as the coefficient of determination from ordinary least squares regression of observed expression on out-of-fold GReX predictions with an intercept. We considered this conventional metric but used its weighted version as the primary evaluation metric to account for heteroscedasticity in normalized pseudobulk expression across individuals. Ignoring covariate and genetic effects, we applied sctransform (Hafemeister and Satija, 2019) to construct inverse-variance weights as

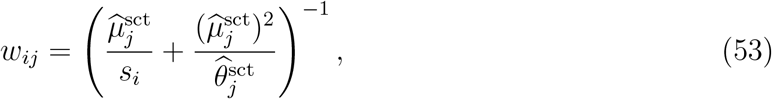

where 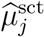 and 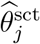 denote the gene-specific mean and dispersion parameters estimated using sctransform, respectively. These evaluation weights were defined independently of any prediction method to ensure a fair comparison. We report weighted and unweighted predictive *R*^2^ as the coefficients of determination from weighted least squares and ordinary least squares regression, respectively. A gene was considered imputable in a given cell type if its weighted predictive *R*^2^ *>* 0.01.

After completing the cross-validation evaluation, we refitted the final GReX prediction models using all available individuals within each cell type. For imputable cell type–gene pairs, the GReX prediction weights from the final fitted models were used in subsequent TWAS analyses.

To assess the contribution of cross-gene correlation to GReX prediction, we performed an ablation analysis. When fitting the joint model in Equation (38), we replaced the estimated cross-gene correlation matrix 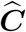 with the identity matrix **I** while keeping all other model components unchanged. We evaluated GReX prediction using the same cross-validation procedure. The ablation analysis was performed within modules containing MOD-scTWAS-only imputable genes and, separately, across all analyzed genes in four cell types with different numbers of cells.

### 4.8 TWAS Analysis of UK Biobank Hematological Traits

We applied the MOD-scTWAS and scTWAS GReX prediction models for imputable genes in each cell type to 29 UK Biobank (UKB) quantitative hematological traits. The selected traits were the same as those analyzed in the scTWAS paper. GWAS summary statistics were obtained from the Pan-UK Biobank resource, and European-ancestry association results were used in the analysis.

Following the FUSION strategy (Gusev et al., 2016), we performed summary-statistics-based TWAS. GWAS summary statistics were harmonized with the OneK1K genotype data by genomic position and allele orientation, with the signs of GWAS *z* scores reversed when necessary. SNPs unavailable in the GWAS summary statistics were excluded from the TWAS calculation. For each gene, let ***w*** and ***z*** denote the vectors of GReX prediction weights and GWAS *z* scores, respectively. The TWAS statistic was calculated as

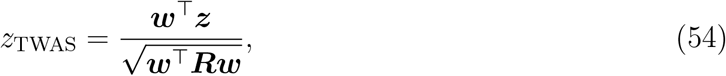

where ***R*** denotes the reference LD matrix for the corresponding cis-SNPs, estimated using genotype data from all 980 OneK1K individuals. TWAS *p* values were calculated from the two-sided tail probabilities of the standard normal distribution. For each trait and cell type, let *m* denote the number of imputable genes for the corresponding method. We used 0.05*/m* as the Bonferroni-corrected threshold for TWAS significance.

## Availability of data and materials

The OneK1K single-cell RNA-seq and genotype data are available through the Gene Expression Omnibus under accession GSE196830. The analyses in this study used raw gene expression counts, imputed genotype data, and covariate files provided by the study authors.

## Supplementary Materials

**Figure S1:**
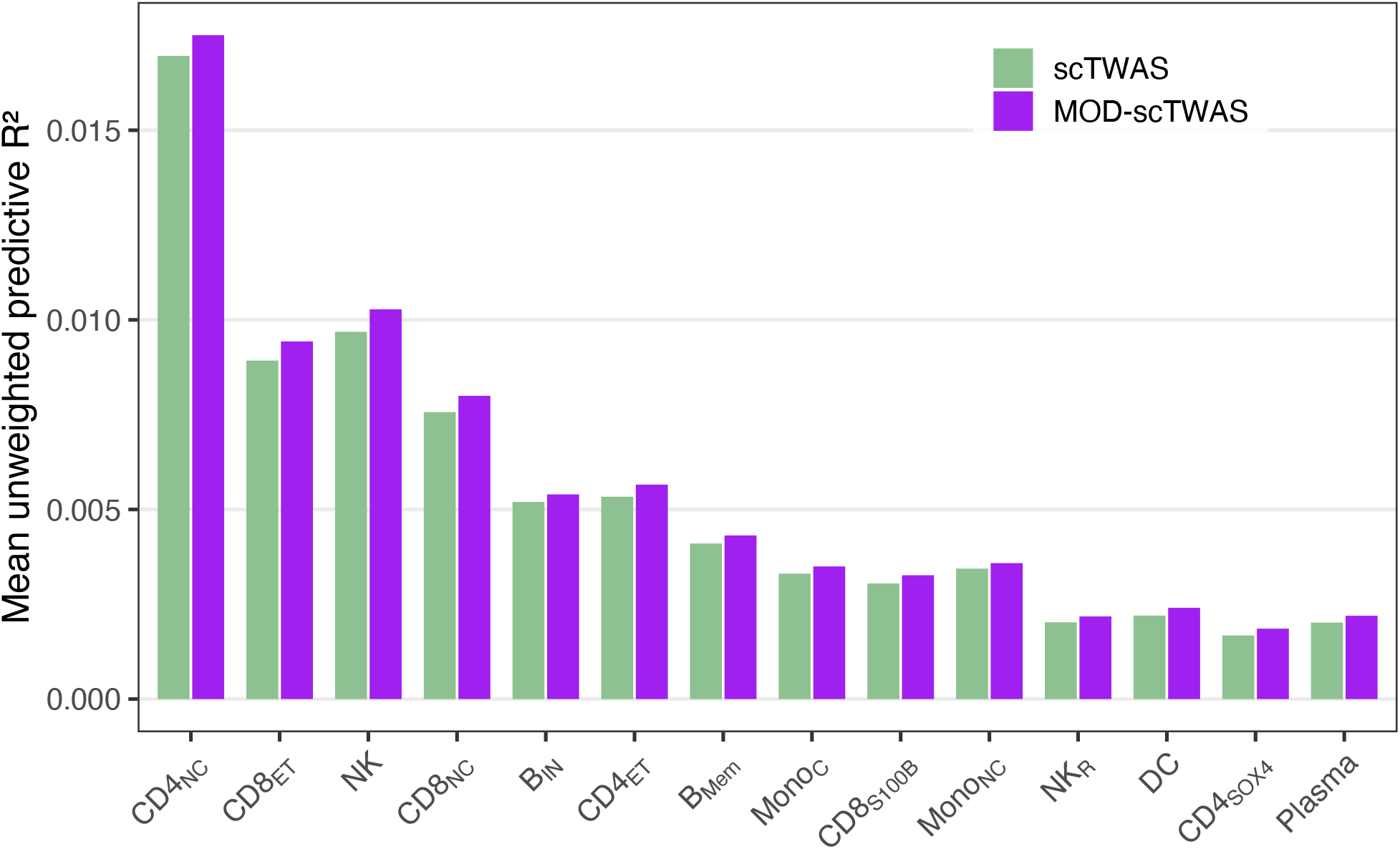
Evaluation of GReX prediction accuracy using unweighted predictive *R*^2^. Mean unweighted predictive *R*^2^ values of scTWAS and MOD-scTWAS across 14 peripheral blood cell types are shown.

**Figure S2:**
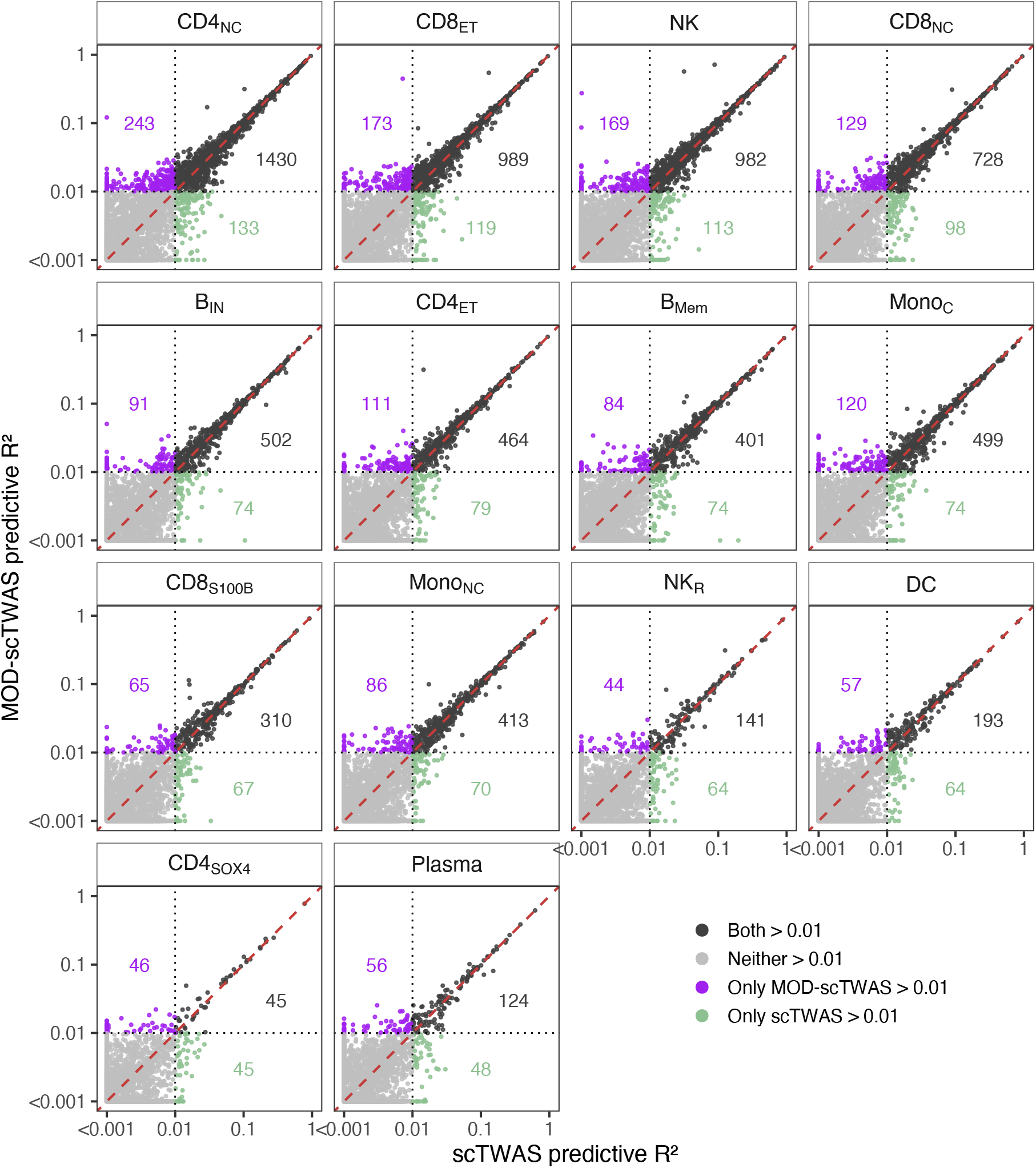
Comparison of gene-level GReX prediction accuracy between scTWAS and MOD-scTWAS. Weighted predictive *R*^2^ values obtained with scTWAS and MOD-scTWAS are compared for individual genes across 14 peripheral blood cell types. Dashed lines indicate the threshold of weighted predictive *R*^2^ *>* 0.01 used to define imputable genes; the numbers indicate the numbers of genes classified as imputable only by scTWAS, only by MOD-scTWAS, or by both methods, respectively.

**Figure S3:**
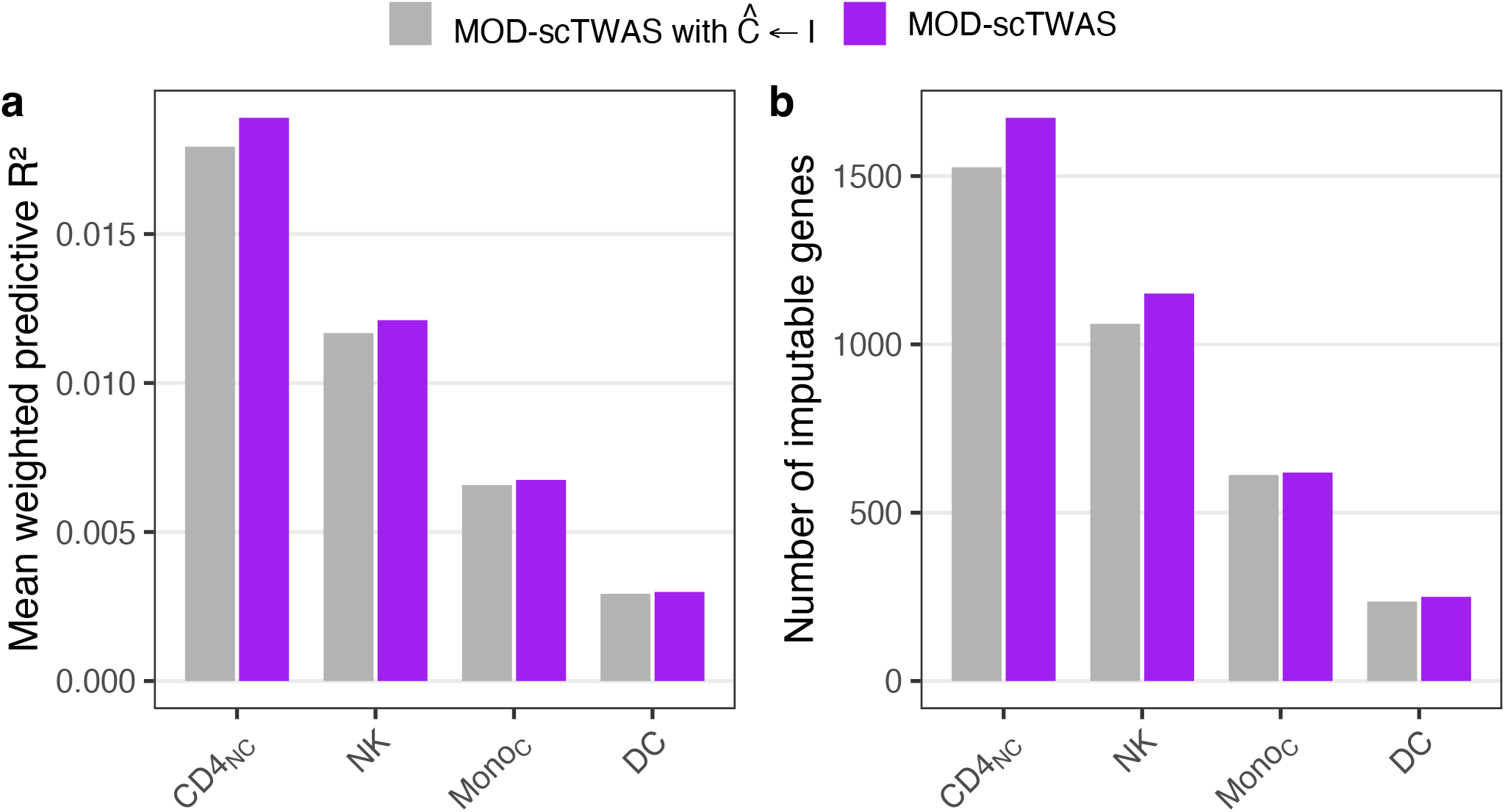
Results of the ablation analysis of cross-gene correlation structure. The full MOD-scTWAS method and MOD-scTWAS with the estimated cross-gene correlation matrix replaced by the identity matrix are compared across four representative cell types in terms of (**a**) mean weighted predictive *R*^2^ and (**b**) the number of imputable genes. Imputable genes are defined as genes with weighted predictive *R*^2^ *>* 0.01.

## Notes

### Competing Interest Statement

The authors have declared no competing interest.

### Summary of Updates

An author affiliation has been corrected to accurately reflect the institutional affiliation of the author. No changes were made to the scientific content of the manuscript.

### Funding

- National Natural Science Foundation of China: T2322017

